# Alignment-Free Guided Design of a Pan-Orthoflavivirus RT-qPCR Assay

**DOI:** 10.64898/2026.03.17.712358

**Authors:** Khanate Sayasit, Chutikarn Chaimayo, Warinya Nuwong, Tharathip Boondouylan, Nattaya Tanliang, Intawat Nookaew, Navin Horthongkham

## Abstract

The co-circulation and rapid expansion of the genus *Orthoflavivirus*, including dengue virus (DENV), Zika (ZIKV), and Japanese encephalitis virus (JEV), pose significant global health challenges. Developing inclusive pan-genus molecular diagnostics is hindered by high nucleotide divergence (>25%–30%) and the computational limitations of traditional multiple sequence alignment in detecting conserved motifs across large datasets. To overcome these limitations, we developed a systematic alignment-free design pipeline that uses rigorous k-mer analysis and compacted De Bruijn graphs. We analyzed 11,846 RefSeq viral genomes to identify phylogenetically conserved, functionally relevant signatures within the *Orthoflavivirus* genus as a case study. The pipeline identified a conserved 600-bp region within the non-structural protein 5 gene, facilitating the design of a broad-spectrum TaqMan RT-qPCR assay. Analytical validation against standard reference strains demonstrated a limit of detection of 1–10 copies/µL for DENV1–4, ZIKV, and JEV, with no cross-reactivity against non-target pathogens. In a clinical evaluation of archived samples, the assay achieved 97.33% overall accuracy. It demonstrated 100% sensitivity and specificity for DENV serotypes, yielding significantly earlier cycle threshold (Ct) values compared to a standard commercial kit, while ZIKV detection showed 100% specificity with 71.43% sensitivity. This study validates an alignment-free, k-mer guided approach for uncovering conserved diagnostic targets in highly variable viral genera. The resulting assay offers a robust tool for frontline surveillance, and the computational framework provides a scalable solution for future pandemic preparedness.

## Introduction

Arthropod-borne viruses (arboviruses) have been a major and expanding global health threat. Members of the genus *Orthoflavivirus*, including dengue virus (DENV), Zika virus (ZIKV), Japanese encephalitis virus (JEV), West Nile virus (WNV), and yellow fever virus (YFV), are responsible for recurrent epidemics, severe clinical manifestations, and substantial socioeconomic burden across tropical and subtropical regions^1^. In 2024, more than 14 million dengue cases and over 10,000 dengue-related deaths were reported globally, underscoring the escalating worldwide burden of this arboviral disease^2^. Asia, particularly Southeast Asia, bears a disproportionate share of the global dengue burden^3,4^. Co-circulation of multiple *Orthoflavivirus* species, rapid geographic expansion, and climate-driven changes in vector distribution have increased outbreak frequency and risk of large-scale epidemics. Climate warming is expanding the geographic range of *Aedes aegypti* (a primary vector of DENV and ZIKV, and the urban-cycle vector of YFV) into higher latitudes and altitudes, while *Culex* species (the mosquito vector for JEV and WNV) responses remain variable. Such expansion threatens to extend flaviviral transmission zones beyond historically endemic tropical areas into temperate regions. In October 2024, the World Health Organization launched the Global Strategic Preparedness, Readiness and Response Plan for dengue and other *Aedes*-borne arboviruses, emphasizing diagnostic innovation as a critical priority for surveillance and outbreak response^5–8^.

Timely and accurate detection is essential for effective surveillance and clinical management of co-endemic *Orthoflavivirus* infections; however, current diagnostic modalities have significant limitations. Serological assays are inexpensive and scalable but suffer from extensive cross-reactivity among flaviviruses, particularly IgG responses in populations with prior exposure or sequential infections, where secondary immune responses confound case attribution. Neutralization testing remains the gold standard for serological differentiation but is impractical for routine surveillance due to biosafety and technical constraints^9^. While whole-genome next-generation sequencing provides unbiased detection and genomic information, it remains costly, slow, and technically demanding for routine frontline screening for clinical diagnosis in low- and middle-income settings. In practice, real-time reverse transcription quantitative PCR (RT-qPCR) has been the primary tool for acute-phase diagnosis and high-throughput surveillance. However, designing pan-*Orthoflavivirus* RT-qPCR assays that retain broad inclusivity across >25%–30% nucleotide-level divergence among distant species while avoiding off-target amplification represents a non-trivial challenge^10–12^. Standard design approaches often fail to simultaneously achieve inclusivity across genetic diversity, specificity against non-target sequences, and robustness against emerging variants not represented in reference databases. Broadly inclusive, genus-level assays that capture phylogenetic diversity while enabling subsequent species- or serotype-level characterization are particularly attractive for surveillance networks in co-endemic regions^11^.

Traditional assay development workflows rely on multiple sequence alignment (MSA) of reference genomes followed by heuristic identification of “conserved” regions for primer–probe placement. This approach carries intrinsic limitations. Even though modern MSA algorithms are computationally robust, the fundamental challenge is distinguishing genuine phylogenetically conserved features from alignment artifacts, especially in regions with competing sequence motifs or low information content. Moreover, generating and maintaining high-quality MSAs becomes increasingly challenging for ultra-large and/or taxonomically broad datasets as sequence databases expand. In practice, conserved-region identification is often biased toward well-studied reference strains (e.g., classical DENV1 Thailand strain, West Nile NY99), under-representing emerging lineages such as dengue genotype V (spreading across Brazil and South America as of 2024), recent Asian Zika genotype variants, or understudied tick-borne species (Powassan, Langat viruses) with limited representation. Together, these factors risk producing nominally “pan-genus” assays with hidden blind spots that show performance decay as novel variants accumulate^13–18^. Alignment-free (AF) methods using k-mer analysis offer scalable alternatives for large-scale genomic data analysis^19,20^. However, researchers have employed AF approaches for whole gene and genome comparison by generating global similarity metrics or phylogenetic signals without consideration of local k-mer variation for designable primer-probe targets^21–24^.

Thus, in this study, we developed a systematic pipeline for the identification of conserved regions, based on AF and k-mer–guided design for high-throughput multiplex RT-qPCR that targets the genus *Orthoflavivirus*. Our approach couples AF k-mer discovery with feature-aware genomic localization and standardized region selection based on i) average common feature (ACF) metric, which is used to identify k-mer length, yielding specific for *Orthoflavivirus* genus (identifying k=19 for 51 orthoflaviviral genomes); ii) compacted de Bruijn graph analysis, which is used to identify conserved k-mer “seeds” that are anchored to functional genomic feature annotations, which ensured primer-binding sites reside within genuinely phylogenetically conserved, functionally relevant regions rather than low-information segments that appear conserved due to alignment artifacts; and iii) local MSA is applied only on the identified conserved region (∼600 bps), down streaming a design aid, avoiding the brittleness of global MSA while still leveraging alignment information where it is reliable. We experimentally validated the designed primer-probe set using standard reference virus samples, and we applied the designed primer-probe set to clinical samples and compared the results with those of commercial kits.

## Materials and Methods

### Data acquisition and preprocessing

The April 2025 FASTA nucleotide files for 18,678 representative viral genomes were downloaded from the NCBI RefSeq database via NCBI Virus, a portal for curated viral sequences and metadata^25^. Taxonomic annotations according to corresponding Taxonomic Identifier (TaxID) were generated with Taxonkit version 0.20.0^26^ using the NCBI Taxonomy dump (September 2025). Segmentation status (segmented vs non-segmented) was annotated using the ViralHostDB database^27^. General feature format 3 (GFF3) annotation files were downloaded for further annotation during block calling.

To reflect biological reality, each genome was represented by a single FASTA file in <taxid>.fasta convention. The downloaded genome records were first filtered by whether they were non-segmented or segmented viruses. Non-segmented genomes were filtered by the keywords in the genome description, and only those with the keywords “complete genome,” “complete sequence,” and “genome” were included. If genomes shared the same taxid, they were assigned an accession number on the filename in the format <taxid_accession>.fasta format, otherwise the filename was labeled as <taxid>.fasta. For segmented viruses, genomes with “partial” in the description, as well as viruses with only one record per TaxID, were discarded. Additionally, genomes that survived the keyword filtering step were kept, resulting in multi-FASTA files used in downstream analyses.

### Alignment-free phylogeny for clade visualization (Mash-style transformed Jaccard index)

To provide a fast overview of clade structure across *Flaviviridae*, we computed a pairwise distance matrix from preprocessed FASTA genomes using Mash distance, which was transformed from Jaccard index to estimate the mutation rate per site between 2 sequences as implemented in the k-mer-length iterative selection for unbiased ecophylogenomics (KITSUNE) dmatrix module. Briefly, genomes were converted to k-mer sets, the exact Jaccard similarity J was computed between k-mer presence/absence (PA) profiles, and distances were obtained via the Mash transform D = (-1/k)(ln(2J/1+J). The resulting distance matrix was used to infer a neighbor-joining tree, which can be used to cluster and visualize clades, not for deep phylogenetic inference^21^. We used a fixed value of 11 for k to avoid parameter overfitting and to remain consistent with KITSUNE’s viral benchmarking, which reported k = 11 as an empirically optimal and subsampling-robust choice for viral comparisons. Conceptually, this value balances 2 competing failure modes: (i) when k is too small, unrelated genomes can share many short k-mers by chance, compressing distances and obscuring clade separation; (ii) when k is too large, overlap between divergent genomes becomes sparse, causing distances to saturate and become unstable. Using k=11 maintains informative overlap for inter-genus comparisons while preserving separation of major clades^28^. The tree was rooted using *Pestivirus* as an out-group to visualize in-group/out-group separation. Tips and clades were colored by genus (*Orthoflavivirus*, *Hepacivirus*, *Pegivirus*, *Pestivirus*, and unclassified *Flaviviridae*) to facilitate qualitative assessment of expected genus-level grouping. Bootstrap support values were not reported because this neighbor-joining tree is used for descriptive visualization rather than inferential phylogenetic conclusions; downstream analyses focus on conserved-window discovery and primer ability rather than resolving deep branching order^29^.

### ACF-guided K Selection and Shortest Unique k-mer Plots (KITSUNE)

We used KITSUNE to guide k selection with the ACF, defined as the mean pairwise intersection size of canonical k-mer sets at length k (reverse-complement canonicalized). KITSUNE ACF can be used to find the maximum informative feature length of k-mers for AF feature frequency profile phylogenomic analysis^28^, and can also identify the shortest unique k-mer (SUK) length for diagnostics with the following method: for a given query genome, sweep k and count how many reference genomes (excluding the query itself) share ≥1 exact k-mer with the query; the SUK is the smallest k for which this count is zero, indicating the presence of a species-specific signature in the database^29^. Moreover, we can infer the length at which a degree of shared k-mers is present.

We performed k-mer counting with Jellyfish version 2.3.1, a fast and memory-efficient tool to count k-mer^30^. In KITSUNE, the “--fast option directs Jellyfish to count all k-mers (no frequency filter), whereas the default run applies Jellyfish’s high-frequency k-mer filter (a two-pass procedure).

We used KITSUNE ACF with the --fast option, so that we included all k-mers present in a genome, to query each *Flaviviridae* genome against the cleaned RefSeq reference set. We then subset the results to the genus *Orthoflavivirus* and plotted ACF vs. k to assess within-group commonness. Counts were reverse-complement canonicalized, and k-mers overlapping non-ACGT characters were ignored.

### Analysis of k-mer Intersection

For each k, KITSUNE ACF recorded every query-reference pair, whether ≥1 exact canonical k-mer was shared. We aggregated these per-pair records into per-k summaries. To reduce storage and complexity, we did not retain the identities of the intersecting k-mers at the global reference level. This is sufficient for all analyses that rely on PA. For downstream conserved-block discovery at k, we performed a separate, targeted re-count within the *Flaviviridae/Orthoflavivirus* set that does capture k-mer identities and supports block merging. If weighted overlaps were required, they were reconstructed from the released FASTA snapshots.

### Presence-absence matrix of k-mers

Using the k-mer length of 19 from the ACF step, genomes were decomposed into canonical 19-mers. We first precomputed Jellyfish k-mer count databases for the full viral collection and *the Orthoflavivirus* subset using Jellyfish count. We then used Jellyfish query/dump to compute exact k-mer membership against the prebuilt universe databases. This method yields a binary PA matrix (columns = 51 *Orthoflavivirus* genomes; rows = distinct universe-referenced 19-mers observed in the *Orthoflavivirus* set). From the PA matrix M, we quantified within-genus conservation for each k-mer as a *present_in* number (the number of *Orthoflavivirus* genomes containing the k-mer) and its fraction = present_in / N. We retained the list of genomes in which each k-mer occurred for downstream QC.

### Compacted De Bruijn graph construction

We used De Bruijn graphs to encode exact (k-1)-base overlaps between k-mers, where each edge is a k-mer and connects 2 nodes that are its (k-1)-prefix and (k-1)-suffix. Long, perfectly consistent stretches of sequence appear in the graph as non-branching paths. By merging each maximal non-branching path into a single node (a unitig), the graph is compacted without loss of sequence information^31^. To obtain compacted De Bruijn graphs and unitigs, we used Bifrost version 1.3.5, a highly parallel construction and indexing of colored and compacted De Bruijn graphs that implements a highly parallel algorithm for direct construction and indexing of colored compacted De Bruijn graphs^32^. The unitigs that were derived from k-mers shared by at least 3 genomes (high-confidence unitigs) were retained for further steps.

### Exact mapping and feature annotation

We mapped the high-confidence unitigs to genomic coordinates by aligning them to a prebuilt Bowtie index of the orthoflaviviral genomes in exhaustive, exact-match mode, reporting all zero-mismatch hits as sequence alignment/map (SAM) records. Bowtie version 1.0.0 was chosen for its speed and strict mismatch control for short sequences^33^. SAM alignments were converted to coordinate-sorted, compressed binary version of SAM files (BAM files) with SAMtools version 1.16.1 (view, sort) and then to browser extensible data (BED) intervals (0-based, half-open) with BEDtools version 2.30.0^34,35^, yielding genomic coordinates for each unitig placement. To link these coordinates to a functional context, we intersected the unitig BED intervals with per-genome GFF3 annotation using BEDtools intersect in strand-agnostic mode, annotating each hit with any overlapping feature.

### Conserved-window selection and scoring

We used a window length of 300 bp, treating this as a design window rather than the final amplicon. This length is approximately 2 to 4 times a typical optimal TaqMan RT-qPCR amplicon (∼70–150 bp), which is generally considered the maximum limit to ensure high efficiency and accurate quantification. It also provides enough local context to place primers and a probe, to tolerate modest sequence variation and small indels, and to accommodate differences in genome length, while still offering fine-scale resolution along the approximate 10–11 kb *Orthoflavivirus* genomes. Features shorter than 300 bp were taken in full as their conserved windows. For longer features, we generated candidate window starts at all unique unitigs hit boundaries within the feature (distinct hit starts, hit ends, and the feature start). For each candidate start, a 300 bp window was placed and, if necessary, shifted so that it remained entirely within the feature bounds.

For every window, we computed 2 scores: i) the summation number of bp of the unitigs mapped on the individual 300 bp windows of the orthoflaviviral genomes, and ii) the number of orthoflaviviral genomes that the unitigs mapped on the individual 300 bp windows. Candidate windows were therefore ranked primarily by the score for further selection of conserved region(s) that were used in the next step.

### Sequence extraction and multiple sequence alignment

For selected high-scoring regions, we extracted the corresponding nucleotide sequences using a 600 bp design region spanning the peak interval and its immediate flanks. This extraction length provides sufficient local context for primer-probe placement while accommodating genome-length variation and small boundary shifts across genomes. Extracted regions were written in a single multi-FASTA file containing sequences from all viruses and multiple sequences were aligned with MAFFT version 7.505 using L-INS-i algorithm, a high-accuracy method using local pairwise alignment.

### Consensus-guided primer design, degeneracy evaluation, and specificity assessment

MSAs of candidate conserved windows were manually inspected for location of conserved identity blocks in Geneious Prime version 2026.0.1 to guide primer–probe placement. Because sequence diversity necessitated the use of degenerate oligonucleotides, IUPAC ambiguity codes were introduced conservatively, and total degeneracy was capped at ≤100 variants per oligo to maintain effective working concentration. A ≥50% per-position base frequency threshold was used to define the consensus base; positions below this threshold were encoded with IUPAC ambiguity codes or avoided when selecting primer-probe binding sites to limit excessive degeneracy.

Primer–probe design rules were applied to optimize amplification and detection efficiency. Forward primer placement was prioritized because mismatches near the 3′ end (defined here as the last 5 bases) can substantially reduce RT-qPCR efficiency; therefore, we preferentially selected regions with contiguous conserved bases and minimized degeneracy toward the 3′ end. In addition, forward primers were constrained to avoid >4 consecutive G residues and to limit G/C content near the 3′ terminus. Probe placement required a conservation level comparable to the forward primer, as mismatches near the probe center can markedly reduce hybridization and cleavage efficiency^36^. Additional probe constraints included avoiding a 5′ G residue, avoiding >4 consecutive G residues, avoiding ≥6 consecutive A residues, and avoiding G at the second position from the 5′ end.

Because reverse transcription is primed by the reverse primer in this one-step assay, limited mismatches may be more tolerated during the RT step with a Moloney murine leukemia virus-derived reverse transcriptase; however, mismatches at the extreme 3′ end were still minimized because both primers must support efficient PCR amplification during cycling^37^. Probe melting temperature (T_m_) was designed to be higher than primer T_m_, and primer T_m_ values were matched as closely as possible. Secondary structures were assessed using predicted hairpin melting temperatures and ΔG values; candidates with hairpins or dimers predicted to be stable, in other words, a strongly negative ΔG, and those with a structure T_m_ approaching the annealing temperature (55 °C), were discarded. All degenerate primers and probes were expanded into explicit sequence variants and evaluated in IDT OligoAnalyzer in batch mode using reaction-relevant oligo and salt parameters^38^. We then assessed the specificity of the designed primers and probe using BLASTN against the NCBI non-redundant nucleotide database, mimicking in silico PCR, and allowing ≤2 mismatches based on primers and probe sequences to approximate assay behavior.

### Standard virus reference strains and RNA extraction

Reference strains of medically important arboviruses were used to evaluate the analytical specificity of the assay. The standard panel included DENV serotypes 1–4 (DENV1–4), ZIKV, chikungunya virus (CHIKV), JEV, WNV, and YFV. Reference viruses were obtained from the American Type Culture Collection (ATCC), including DENV1 strain Hawaii (ATCC VR-1856), DENV2 strain TH-36 (ATCC VR-1810), DENV3 strain H87 (ATCC VR-3380), DENV4 H241 (ATCC VR-1490), and ZIKV strain PRVABC59 (ATCC VR-1843). JEV strain P3 was obtained from the Faculty of Tropical Medicine, Mahidol University, Thailand. Nucleic acid controls for WNV strain New York-99 (MBC069-R) and YFV strain 17D (MBC100-R) were obtained from VIRCELL Molecular. All viruses except WNV and YFV were cultured and subjected to lysis before RNA extraction. RNA was extracted from 200 µL of supernatant using the MagLEAD 12gC automated extraction platform (Precision System Science) following the manufacturer’s protocol and eluted in 100 µL of RNase-free water. Extracted RNA was either used immediately or stored at −80 °C for short-term use.

### Evaluation of analytical specificity using an arboviral standard viral panel and cross-reactivity testing

To assess the potential detection of the designed primers and probe, we tested RNA extracted from each reference virus using the developed real-time RT-qPCR assay. Real-time RT-qPCR was performed in a total volume of 20 µL using qScript® XLT One-Step RT-qPCR ToughMix®, Low ROX™ master mix (Quantabio). Each reaction contained 2x master mix, forward primer (500 nM), reverse primer (500 nM), probe (125 nM), and 5 µL of template. Nuclease-free water was added to reach the final volume. Reactions were run on a QuantStudio™ 7 Pro Real-Time PCR System (Thermo Fisher Scientific) under the following cycling conditions: cDNA synthesis at 50 °C for 10 mins, initial activation at 95 °C for 1 min, followed by 45 cycles of denaturation at 95 °C for 10 seconds and combined annealing/extension at 55 °C for 60 seconds with fluorescence acquisition at the end of each cycle. Baseline and threshold values were automatically calculated by the instrument software and manually adjusted if necessary. Cross-reactivity testing was performed to assess the analytical specificity of the designed primer–probe set. The assay was evaluated in a single-reaction format using nucleic acids from the following non-target pathogens: Influenza A virus H1N1 strainA/PR/8/34 (ATCC VR-95), Influenza B virus strain B/Lee/40 (ATCC VR-1535), respiratory syncytial virus (RSV) strain A2 (ATCC VR-1540), CHIKV strain ROSS (Department of Medical Sciences, Ministry of Public Health, Thailand), enterovirus A71 (EV-A71) strain H (ATCC VR-1432), *Orientia tsutsugamushi* strain KARP (BioSample accession SAMN51290297), and *Rickettsia typhi* strain SI-typh0423 (BioSample accession SAMN51290298). Each pathogen was tested at approximately 2 × 10^2^ copies/µL, except CHIKV, EV-A71, and RSV, which were tested at 2 × 10^4^ copies/µL.

### Determination of copy number

A kit-provided positive-control template (2 × 10^5^ copies/µL) was serially diluted 10-fold in template preparation buffer to generate standards of 2 × 10^4^, 2 × 10^3^, 2 × 10^2^, 2 × 10^1^, and 2 copies/µL. For standard-curve generation, 5 µL of each dilution was added to 15 µL reaction master mix (final volume, 20 µL) containing the corresponding primer–probe set, corresponding to 10–10^5^ copies per reaction across the dilution series. Standard curves were constructed by linear regression of Ct values against log10 (copies per reaction). Standard-curve reactions for DENV, ZIKV, and JEV were performed using the SuperScript III One-Step RT-PCR System with Platinum Taq DNA Polymerase (Invitrogen). (See **Supplementary Fig. 1**).

### Limit of detection, repeatability, and reproducibility of the designed primers and probe set

The limit of detection (LOD) and repeatability of the designed primers and probe set were evaluated using extracted RNA of 6 arboviruses reference strains, including DENV1–4, ZIKV, and JEV. LOD, and intra-assay repeatability was assessed in 12 replicates of various concentrations (1 to 1,000 copies/μL). Inter-assay reproducibility was evaluated by 2 technicians over 5 days, with a total of 20 replicates. The real-time RT-qPCR reaction was performed as previously described. The standard deviation (SD) and coefficient of variance (CV) were calculated. The Ct cut-off was calculated from the mean at LOD dilution + 2 SD.

### Pathogen-specific commercial kits used

Commercial assays were performed using the Primerdesign™ genesig advanced kits for ZIKV and DENV (Primerdesign™ Ltd). We performed the assays of the commercial kits, following the manufacturer’s protocols.

### Performance on clinical samples

We validate the sensitivity and specificity of the designed primer-probe set using 150 archived clinical samples with prediagnosed arboviral infection, including 25 plasma samples positive for DENV1, DENV2, and DENV4, 11 positive for DENV3, 14 positive for ZIKV, as well as 50 negative plasma samples. Prior to the designed primer-probe validation, all positive samples were confirmed using the pathogen-specific commercial real-time PCR assay (Primerdesign^TM^ Ltd. genesig® advanced kit), which was used as the standard test. All 50 negative samples were also confirmed to be negative for all flaviviruses. For the real-time analysis, 5 µL of extracted nucleic acids were mixed with 15 µL of mastermix (total volume of 20 µL) using qScript® XLT One-Step RT-qPCR ToughMix® (Low ROX™) (Quantabio). Each reaction contained 2x master mix, forward primer (500 nM), reverse primer (500nM), and 125 nM of probe. The real-time RT-qPCR was performed as previously described. The detail of the archived clinical samples is provided in **(Supplementary Table 1A,1B)**.

### Ethic statement

The study was reviewed and approved by the Institutional Review Board at Mahidol University, Bangkok, Thailand (MU-MOU COA No. 576/2023) Additional clinical samples used for validation were obtained from archived samples from our studies (COA No. SI 604/2023 and COA No. SI 946/2024).

### Statistical analysis

Ct values obtained with the designed primer-probe set were compared with those from the commercial kit. To summarize assay repeatability and the distribution of Ct values, we calculated the SD, as well as the median and interquartile range. We also reported the percentage coefficient of variation (%CV), which summarizes variability in Ct values across replicates. Diagnostic performance was assessed by calculating sensitivity, specificity, positive predictive value, negative predictive value, and overall accuracy, together with 95% confidence intervals (95% CI), using MedCalc’s diagnostic test calculator (https://www.medcalc.org/calc/diagnostic_test.php). Bland–Altman plots were used to evaluate agreement between the 2 assays across the Ct range. Differences in Ct values between the designed primer-probe set and the commercial kit were assessed using a two-sided paired t-test, and Pearson correlation was used to assess the strength of the linear relationship between Ct measurements. Plots were generated in GraphPad Prism 10.

## Results

The workflow of our systematic primer and probe design for pan detection of *Orthoflavivirus* genus is summarized in **Fig. 1A**. The key steps of the workflow are described in the following sections.

**Fig. 1.**
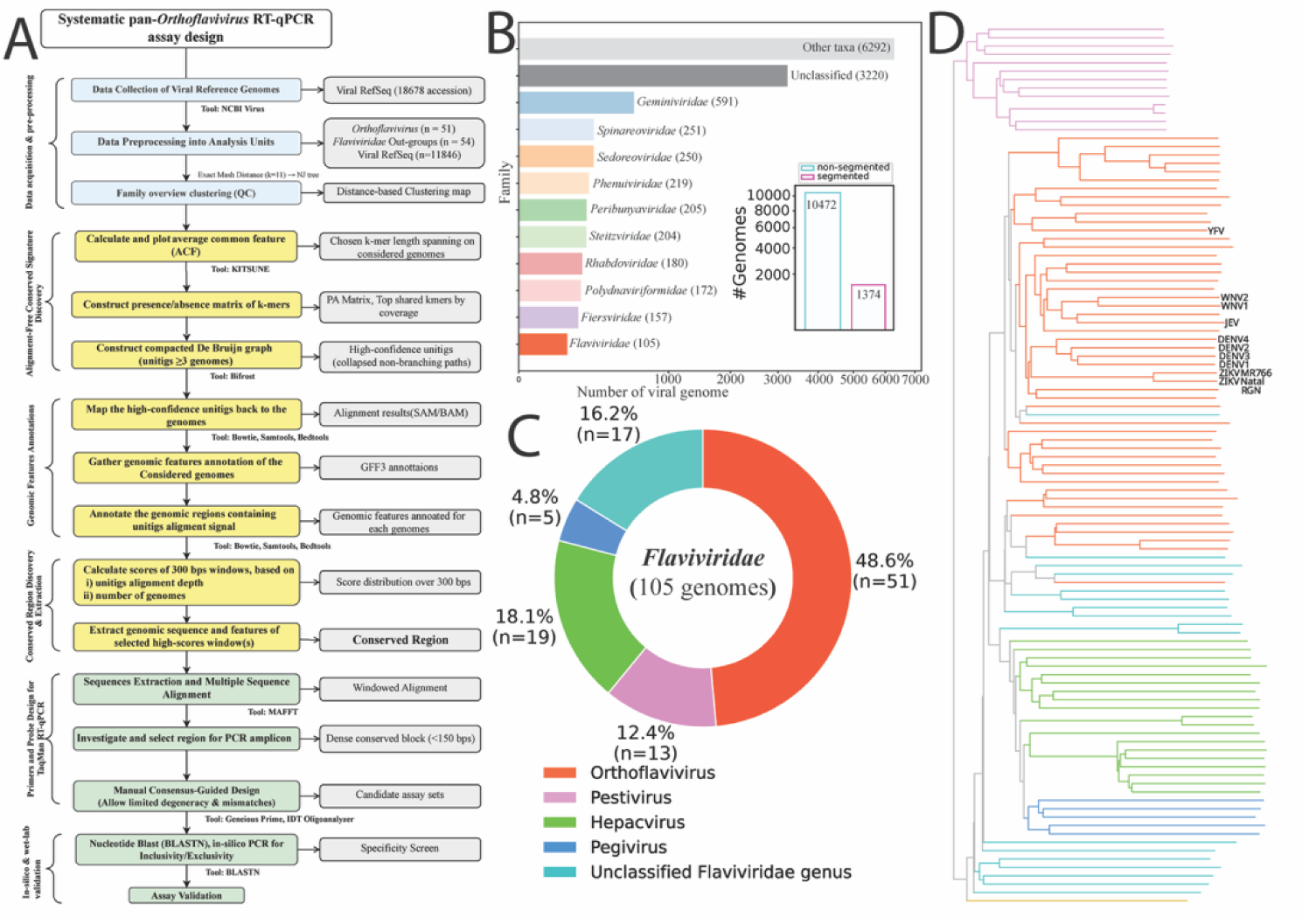
A comprehensive workflow for a pan-*Orthoflavivirus* RT-qPCR assay design, from dataset curation and alignment-free conserved-window discovery for assay validation. **(A)** The workflow is used to (i) verify dataset structure/QC; (ii) localize and rank conserved, feature-anchored windows; and (iii) guide MSA-based manual primer-probe design and specificity screening. Box colors indicate workflow phases; grey boxes denote intermediate artifacts/outputs; yellow boxes mark workflow components introduced in this study. **(B)** Family-level representation of dataset composition. Bars show the top 10 viral families by number of genomes, with Flaviviridae shown additionally for reference (highlighted), even though it falls outside the top 10. The cleaned dataset comprises 11,846 viral genomes, including 10,472 non-segmented and 1,374 segmented genomes. Most genomes fall into other taxa (6,292) and Unclassified (3,220), followed by Geminiviridae (591), Spinareoviridae (251), Sedoreoviridae (250), Phenuiviridae (219), Peribunyaviridae (205), Steitzviridae (204), Rhabdoviridae (180), Polydnaviriformidae (172), Fiersviridae (157), and Flaviviridae (105). **(C)** Genus-level composition of the family Flaviviridae in the RefSeq complete/near-complete genome dataset. *Orthoflavivirus* comprises the largest fraction (48.6%, n = 51), followed by *Hepacivirus* (8.1%, n = 19), unclassified Flaviviridae (16.2%, n = 17), *Pestivirus* (12.4%, n = 13), and *Pegivirus* (4.8%, n = 5). Genera are ranked by the number of complete/near-complete RefSeq genomes. **(D)** An alignment-free distance matrix was computed from preprocessed *Orthoflavivirus* genomes using a Mash-style transformed Jaccard metric (k = 11) and used to infer a neighbor-joining tree (rooted with a *Pestivirus*, orange branch in the bottom). The topology recapitulates expected intra-genus structure and coherent species-level clades, which we used to verify taxonomic assignments and flag outliers prior to conserved-signature analysis. The focused species within this study were labeled with their names on the tree. DENV1–4 = dengue virus serotype 1-4, JEV = Japanese encephalitis virus, WNV1,2 = West Nile virus linage 1,2, ZIKV_MR766 = Zika virus, ZIKV_ Natal_RGN = Zika virus

### Data acquisition and preprocessing: *Viral Genome Dataset Composition and Phylogenetic Validation*

We began by downloading 18,678 accession numbers of viral genomes from NCBI RefSeq. After filtering out non-genome accession numbers, we were left with 15,467 accessions, comprising 10,472 non-segmented viral genomes and 1,374 segmented viral genomes. In total, 11,846 viral genomes were used for further analysis as a reference genome set. The distribution of the top 10 viral families by number of genomes is illustrated in **Fig. 1B** (all genomes and annotations are summarized in **Supplementary Table 2A and 2B)**. Focusing on the *Flaviviridae* viral family (**Fig. 1C**), 51 from 105 genomes were assigned to the *Orthoflavivirus* genus, and another 54 genomes belonged to other genera—*Pestivirus, Hepacvirus, Pegivirus,* and Unclassified— providing both in-group and closely related out-group context for conserved-signature discovery analysis. The AF phylogenomic relationships were reconstructed and illustrated in **Fig. 1D**. The expected clustering of *Orthoflavivir*us genomes was clearly observed in the phylogenomic tree, supporting the taxonomic coherence of the dataset used for downstream analysis. In addition, selected clinically important *Orthoflavivirus* species such as DENV1–4, JEV, WNV lineages 1 and 2, YFV, and 2 ZIKVs, marked on the tip of the phylogenomic tree, were used for experimental validations in the study.

### Alignment-free conserved sequence signature discovery: Identification and graph-based consolidation of conserved k-mer signatures

To obtain our choice of k-mer length that was used as seeds for conserved sequence signature identification, covering the *Orthoflavivir*us genus, we used KITSUNE^28^ to calculate ACF, representing the pairwise genome similarity based on k-mer profiles at a specific k-mer length. The individual *Flaviviridae* genomes were compared against the set of 10,472 reference genomes to determine whether the genomes had common k-mers or not. The number of genomes with k-mer sequences in common with the 105 genomes in *Flaviviridae* was recorded for each k-mer length between 9 and 51 and plotted for comparison (**Fig. 2A**). From the plot, we identified the SUK length of the *Flaviviridae* genomes, highlighting *Orthoflavivirus* queries. Each curve represents a single *Flaviviridae* genome; at each k-mer length it reports the number of genomes in the full viral reference collection (including itself) that share ≥1 canonical k-mer. We found that at a k-mer length of 23, a specific k-mer can be found for the *Orthoflavivirus.* Considering the genomes of standard reference viruses (**Fig. 2B**), the k-mer length 31 onward yielded specific k-mers. From the plots (**Fig. 2A and 2B**), at the cut point of 51 orthoflaviviral genomes, we found that a k-mer length of 19 nt was the shortest k-mer length of the *Orthoflavivirus* genus that contained specific common k-mer(s) among them. In total, 526,787 unique k-mers were identified for the 51 orthoflaviviral genomes. We found that the top 2 k-mers were commonly found in at maximum 20 genomes (**Fig. 2C**). To reduce the number of trivial sliding window duplicates, the unique k-mers were used as the seed for a compacted De Bruijn graph based on sequence overlapping among them to produce unitigs derived from the aggregation of k-mers within the close neighbors. We obtained 4,814 unitigs in total. Only 339 of these unitigs had sequences present in at least 3 orthoflaviviral genomes, and thus were designated high-confidence unitigs (**Fig. 2D**) and used for further steps. The high-confidence unitigs ranged from 19–41 bp with a maximum k-mer support of 23 and a maximum number of unitigs present in the genomes of 20.

**Fig. 2.**
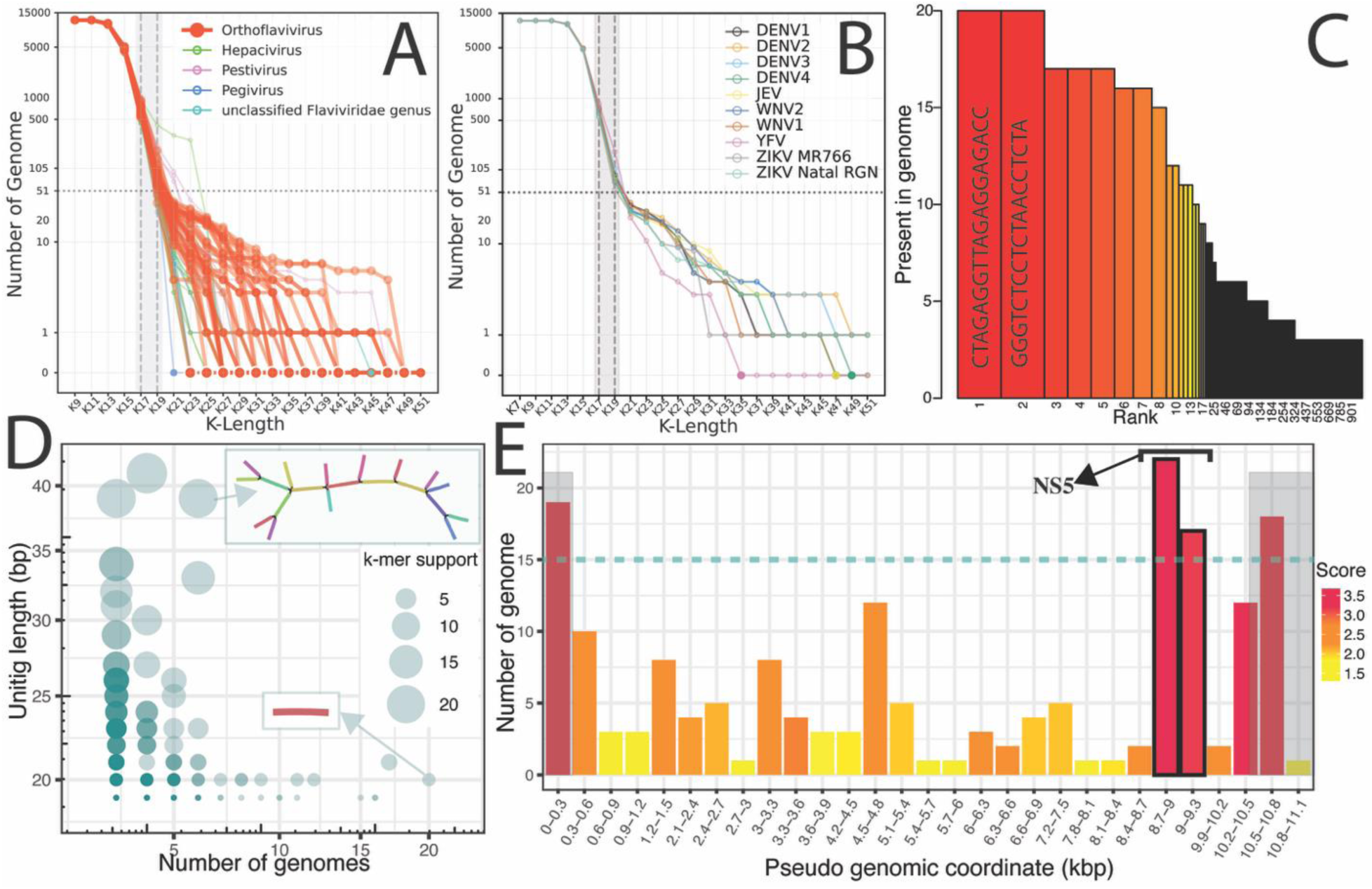
Alignment-free conserved-signature discovery and localization in *Orthoflavivirus* genomes. **(A)** Average common feature shortest-unique k-mer (SUK) curves across k=9–51. Vertical guides highlight the onset of rapid feature loss from K17 and a sharper decline around K19, marking the transition from broadly shared to increasingly specific k-mer features. **(B)** SUK plot for the *Orthoflavivirus* considered genomes—dengue virus (DENV1–4), Japanese encephalitis virus (JEV), West Nile virus (WNV), yellow fever virus (YFV), and Zika virus (ZIKV)—showing the decay in shared k-mer features as k increases. We selected k = 19 for downstream conserved-signature discovery because at k ≥ 21 the number of shared k-mers dropped below the genus-level panel size (n = 51), limiting detection of features conserved across the full panel. **(C)** Genome-presence rank plot of 19-mers within *Orthoflavivirus*. Bars show the number of genomes containing each 19-mer sequence, ranked from highest to lowest; the top 2 highest-presence 19-mers are labeled. **(D)** Unitig summary after compacted de Bruijn graph (cDBG) consolidation of 19-mers filtered to those present in ≥3 genomes. Unitig length is plotted against the number of genomes supporting each unitig. Point size indicates k-mer support (number of distinct 19-mers per unitig). Highly conserved unitigs (supported by more genomes) are generally shorter, whereas longer unitigs are typically supported by fewer genomes. **(E)** Genome-wide conservation hotspot profile along a pseudo-genomic coordinate. Bars indicate the number of genomes contributing conserved hits per window, with color encoding conservation score. Prominent peaks coincide with the NS5 region; terminal repeat/UTR regions are indicated in gray (0–0.3 kbp and 10.5–11.1 kbp). The gray boxes indicate the terminal repeat/UTR regions.

### Conserved region(s) discovery and extraction

The high-confidence unitigs were then mapped back to the 51 orthoflaviviral genomes using exhaustive, exact alignment with Bowtie. The vast majority of unitigs mapped to one or more loci in the reference panel without mismatches, confirming that the compacted De Bruijn graph-derived sequences faithfully represented k-mers present in the underlying genomes. Conservation scores were calculated by aggregating 300-bp windows from orthoflaviviral genomes. The distribution of the scores over pseudo-genomic coordinates is illustrated in **Fig. 2E**. At the cut-off of 15 genomes, 4 300-bp windows within the pseudo-genomic coordinates were identified, 2 of which were 0–0.3 kb and 10.5–10.8 kb untranslated region/terminal repeats based on the genomic annotation results, and were neglected. Therefore, the 2 windows of 8.7–9.0 kbp and 9.0–9.3 kbp were combined as the conserved region for the probe/primer design. Notably, the 600-bp conserved region is located on non-structural protein 5 (NS5).

### Primers and probe design for TaqMan RT-qPCR

The sequences of the identified 600-bp conserved regions of the 51 individual orthoflaviviral genomes were extracted then multiple sequence alignments were performed using MAFFT software. The result is illustrated in **Fig. 3A**. The consensus sequence derived from the multiple sequence alignments has a length of around 930 bp, including some sequence blocks of conservation (forest green color on identity lane, indicating 100% identity). We next carefully inspected the multiple sequence alignment result using Geneious Prime software to find a region that can produce an amplicon size ∼150 bp containing complement-specific target DNA sequences at both 3’ and 5’-end that are conserved across the species to initiate PCR amplification. We found a region in the middle, between positions 380–530 in the alignment, containing a dense block of conserved sequence that could be the target for an RT-qPCR primer-probe design (region between the 2 red vertical lines in **Fig. 3A**)

**Fig. 3.**
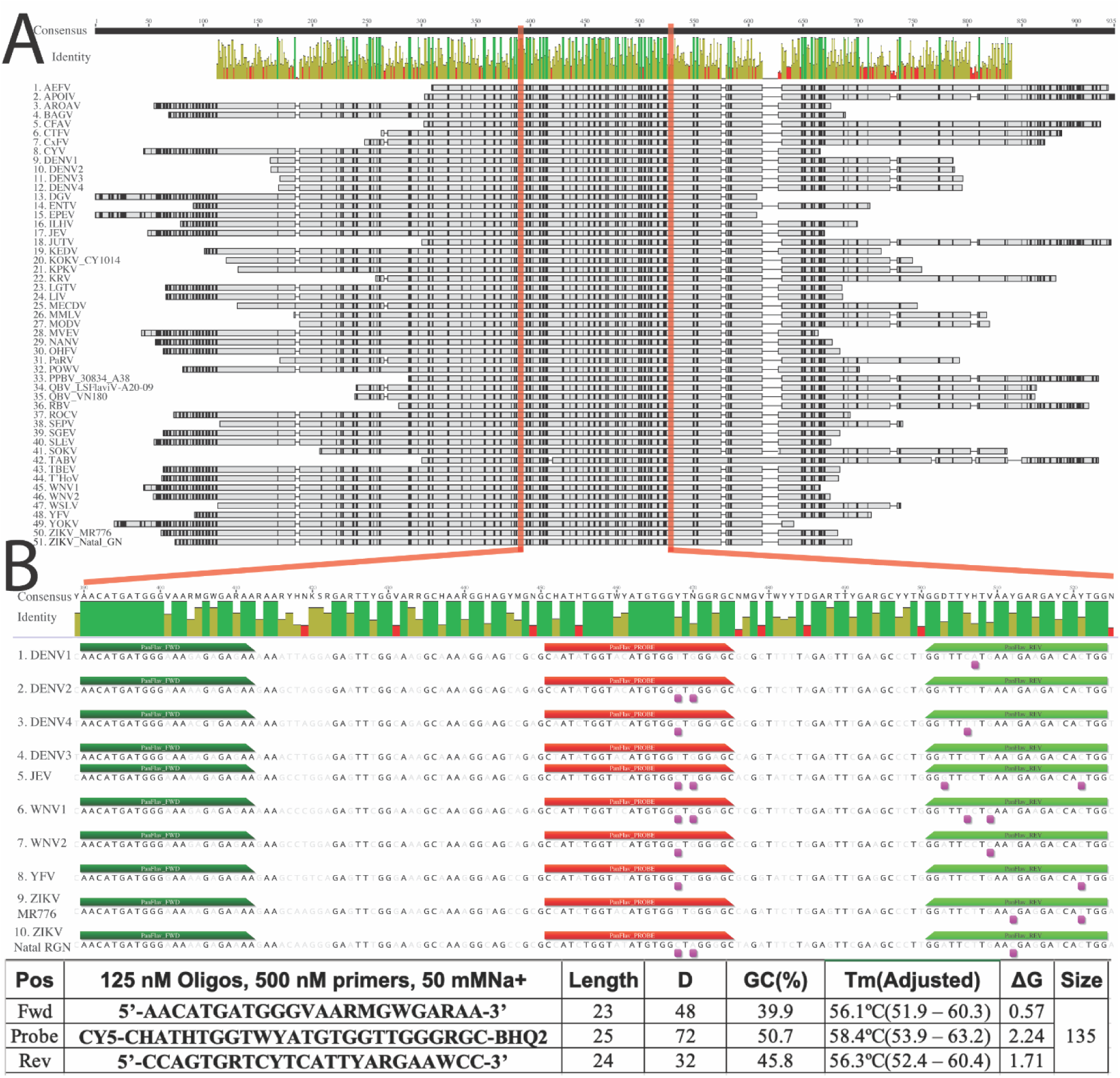
Alignment-free conserved-signature discovery and localization in *Orthoflavivirus* genomes. (A) MAFFT multiple sequence alignment of 600 bps conserved windows from 51 *Orthoflavivirus* genomes, the red lines indicate the start and the end of the amplicon window used to place the primers and probe. The amplicon region shows high per-base identity, indicated by multiple, contiguous, or near-contiguous conserved bases at 100% identity, colored in forest green. (B) The top panel shows MSA of the considered genomes, including dengue virus (DENV1–4), Japanese encephalitis virus (JEV), West Nile virus (WNV1,2), yellow fever virus (YFV), and Zika virus (ZIKV) strains MR766 Natal RGN annotated with primer pair and probe position with mismatches in magenta underneath the genomes. The lower panel shows chemical properties of each oligonucleotide, including probe and primer concentration, Na+ concentration, length, degeneracy, GC content, salt-corrected T_m_, ΔG of the reaction, and product size.

To reduce the complexity of the primer-probe design, we focused only on a subset of 8 medically important orthoflaviviral arboviruses—DENV1–4, ZIKV, JEV, WNV, and YFV. The selected region contains sufficient conserved stretches to place primers and probes while accommodating the observed diversity (**Fig. 3B**). We designed a primer-probe set **(Fig. 3B**, lower panel) that possibly can be used to detect all the 8 of the considered viral species. The set was constrained to avoid mismatches or degeneracy in the last nucleotides of the forward primer and in the central portion of the probe, while permitting limited internal degeneracy or mismatches in the reverse primer when necessary to retain coverage. We observed some sequence variation across different genomes distributed on the complement sequences of the primers and probes; therefore, nucleotide degeneration of the primers and probes was iteratively applied based on the sequence variation context to result in a maximum mismatch of 2 (**Fig. 3B**, top panel) with an appropriate T_m_ and ΔG. The details of the designed primer-probe set are illustrated in **Fig. 3B** (lower panel). The primer-probe set used in the assay comprises a 23-nt forward primer, a 24-nt reverse primer, and a 25-nt hydrolysis probe (5’-Cy5, 3’-BHQ2), yielding a 135 bp amplicon. Across the 3 oligonucleotides, GC content ranged from 39.9%–50.7%, and predicted T_m_ under the specified reaction conditions (125 nM oligos, 500 nM primers, 50 mM Na^+^) were tightly matched (forward, 56.1 °C; reverse, 56.3 °C; probe, 58.4 °C; adjusted ranges shown in the figure). Thermodynamic estimates further indicated favorable hybridization (ΔG = 0.57–2.24), consistent with stable target recognition. We then checked whether the primer-probe set could be used to detect other orthoflaviviruses in the selected region in silico using Geneious Prime software. With 2 mismatches allowed, we found that the primer-probe set can possibly detect 15 Orthoflaviviruses (see **Supplementary Fig. 2A**). Furthermore, we checked the specificity of the primers and probe sequences on the NCBI non-redundant nucleotide database by performing nucleotide blast (BLASTN) with 2 mismatches allowed and found 26466 hits. Almost every hit was a known *Orthoflavivirus*. We deduplicate by TaxIDs, resulting in 42 records, Guapiaçu virus was represented by multiple accessions (TaxID = 2602438), and Panmunjeom flavivirus was represented by accession KY072986.1 (TaxID = 1928710) (See **Supplementary Table 3A, 3B**). Interestingly, these 2 species were flaviviruses newly isolated from mosquitoes^39,40^ that have not been fully taxonomically classified; however, we hypothesize that they are in the *Orthoflavivirus* genus.

We also designed additional primer-probe sets for the 28 *Orthoflavivirus* that could not be detected by the previous assay in silico on the same conserved region. Following the same criteria, 4 additional primer-probe sets will be required in combination to detect all the 51 *Orthoflavivirus* genera (See **Supplementary Fig 2B-2E and Supplementary Table 4A-4D** for details).

### Performance evaluation of the designed primer-probe set for TaqMan RT-qPCR assay

Firstly, we performed a pilot experiment to test the capability of the designed primer-probe set with the standard reference samples of the medically important arbovirus at an arbitrary copy number per microliter, depending on all the available materials. As illustrated in **Fig. 4A**, the designed primer-probe set was able to detect DENV1–4, ZIKV, and JEV within a range from 10–10,000 copies per microliter and was able to detect WNV and YFV at 1,000 copies per microliter (only 1 point was tested due to limited sample material).

**Fig. 4.**
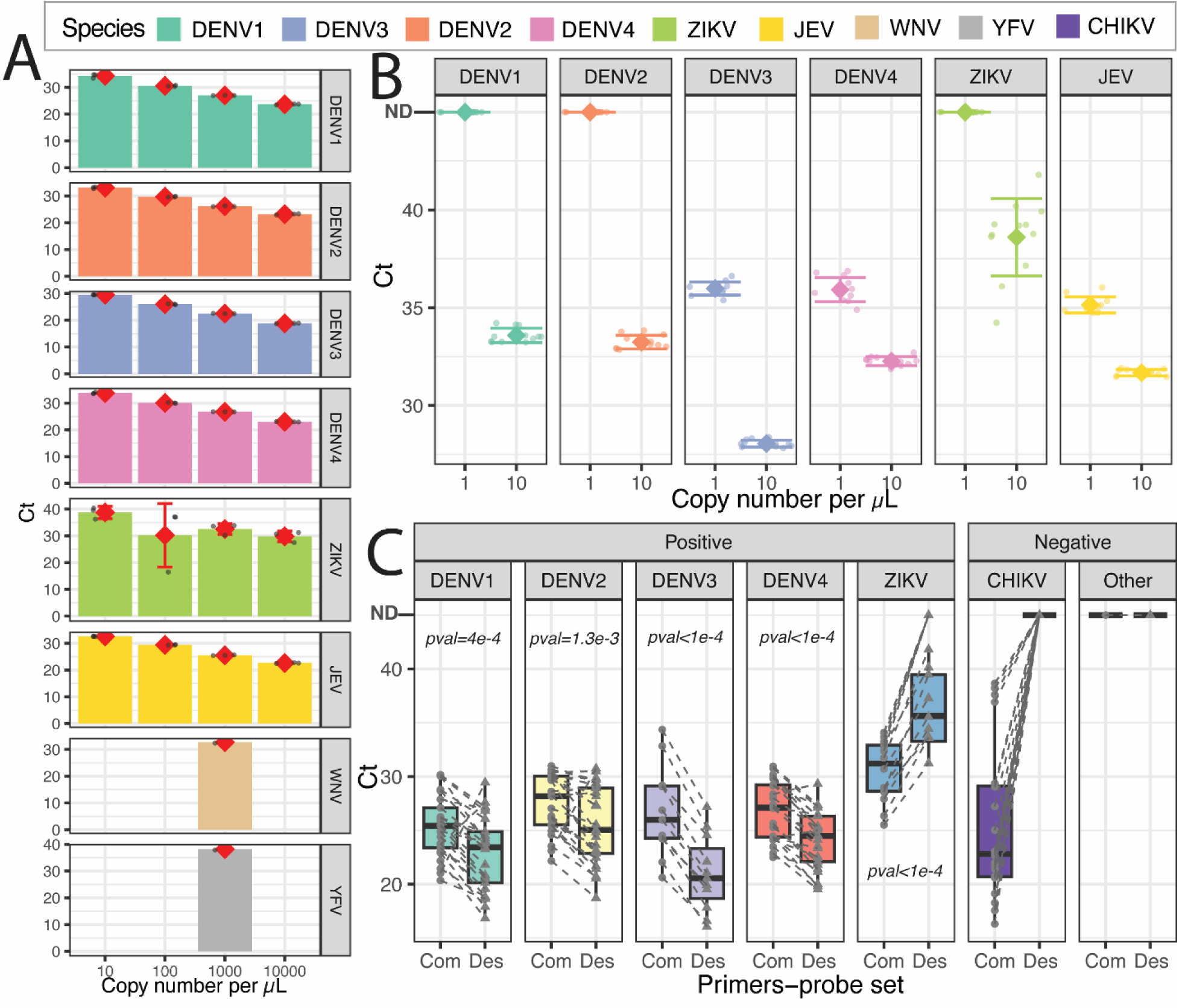
Alignment-free conserved-signature discovery and localization in *Orthoflavivirus* genomes. **(A)** Faceted bar charts show Ct values measured across input copy numbers (copies per µL) for each virus—dengue virus (DENV1–4), Japanese encephalitis virus (JEV), West Nile virus (WNV), yellow fever virus (YFV), and Zika virus (ZIKV). Bars summarize Ct at each concentration, with points indicating individual reactions (technical replicates); red diamonds denote the mean (error bars, where shown, indicate variability across replicates). Across DENV1–4, ZIKV, and JEV (triplicates), Ct values generally decrease as input concentration increases. ZIKV shows greater replicate-to-replicate variation at some concentrations, reflected by wider error bars compared to other targets. WNV and YFV were detected at 1,000 copies per µL; only a single concentration was tested for these viruses due to limited sample material. Together, these results indicate broad target coverage and reliable amplification across a wide input range for the main panel members. **(B)** Limit of detection (LOD) assessment of the designed primer-probe set. Prior to testing clinical specimens, LOD was evaluated using nucleic acids from 6 arboviruses (DENV1–4, ZIKV, and JEV) at 1, 10, 1000 copies per µL, with 12 technical replicates per dilution. The assay detected DENV3, DENV4, and JEV down to 1 copy per µL, whereas DENV1, DENV2, and ZIKV were consistently detected down to 10 copies per µL. Points represent individual reaction; a diamond indicates the mean Ct, and error bars indicate ±SD. ND indicates not detected (did not meet the ≥95% positivity criterion and therefore did not qualify as the LOD). **(C)** Paired comparison of Ct values between assays in clinical samples. Ct values obtained with the commercial kit (Com; circles) and the designed primer–probe set (Des; triangles) are shown for each target, with paired measurements from the same sample connected by dashed lines, for 100 positive samples and 50 positive samples (20 positive with Chikungunya virus [CHIKV] and 30 others). Box plots summarize the distributions (center line, median; box, interquartile range; whiskers, range). Relative to the commercial kit, the designed primer–probe set detected DENV1–4 significantly earlier (lower Ct; *p* = 0.0004 [n = 25], 0.0013 [n=25], <0.0001 [n=11] and <0.0001 [n=25] for DENV1–4, respectively) but detected ZIKV later (higher Ct; *p* < 0.0001 [n =14]). P values were calculated using a two-sided paired t-test on matched Com–Des Ct values for each target. Overall diagnostic performance is reported in **Tables 1,2**.

Before applying the designed primer-probe set on clinical samples, the LOD of primer-probe was determined using the nucleic acids of 6 arboviruses at 1, 10, 100, and 1,000 copies/μL in 12 replicates per dilution. The assay could detect DENV3, DENV4, and JEV at a level of 1 copy/μL and DENV1, DENV2, and ZIKV at 10 copies/μL **(Fig. 4B, Supplementary Table 5)**. The %CV of arbovirus detection was less than 5% in all cases, except for ZIKV, for which it was 5.13% **(Table 1)**.

**Table 1.** Intra-assay repeatability and inter-assay reproducibility of the designed primer-probe set.

| Pathogens | Intra-assay repeatability |  |  |  |  |  |  | Inter-assay reproducibility |  |  |  |  |
| --- | --- | --- | --- | --- | --- | --- | --- | --- | --- | --- | --- | --- |
|  | 1,000 copies/μL |  | 100 copies/μL |  | 10 copies/μL |  | 1 copy/μL | 1,000 copies/μL |  |  | 100 copies/μL |  |
|  | detected/<br>replicates) | Mean<br>(SD, % CV) | detected/<br>replicates | Mean<br>(SD, % CV) | detected/<br>replicates | Mean<br>(SD, % CV) | detected/<br>replicates | Mean<br>(SD, % CV) | detected/<br>replicates | Mean<br>(SD, % CV) | detected/<br>replicates | Mean<br>(SD, % CV) |
| DENV1 | 12/12 | 26.35<br>(0.12, 0.46) | 12/12 | 30.05<br>(0.19, 0.62) | 12/12 | 33.58<br>(0.36, 1.09) | 0/12 | - | 20/20 | 27.33<br>(0.73, 2.65) | 20/20 | 31.09<br>(0.76, 2.45) |
| DENV2 | 12/12 | 26.20<br>(0.30, 1.14) | 12/12 | 29.13<br>(0.15, 0.50) | 12/12 | 33.24<br>(0.34, 1.04) | 1/12 | 42.54<br>(ND, ND) | 20/20 | 26.79<br>(0.54, 2.02) | 20/20 | 30.43<br>(0.66, 2.16) |
| DENV3 | 12/12 | 20.62<br>(0.11, 0.55) | 12/12 | 24.19<br>(0.30, 1.22) | 12/12 | 28.05<br>(0.18, 0.63) | 12/12 | 35.98<br>(0.33, 0.92) | 20/20 | 20.71<br>(0.48, 2.33) | 20/20 | 24.26<br>(0.41, 1.67) |
| DENV4 | 12/12 | 25.18<br>(0.17, 0.68) | 12/12 | 28.49<br>(0.21, 0.74) | 12/12 | 32.26<br>(0.23, 0.71) | 12/12 | 35.92<br>(0.62, 1.72) | 20/20 | 25.65<br>(0.30, 1.15) | 20/20 | 29.15<br>(0.41, 1.39) |
| ZIKV | 12/12 | 32.52<br>(0.30, 0.93) | 12/12 | 35.89<br>(0.29, 0.82) | 12/12 | 38.60<br>(1.98, 5.13) | 3/12 | 40.47<br>(1.89, 4.67) | 20/20 | 33.79<br>(0.62, 1.84) | 20/20 | 37.68<br>(0.63, 1.66) |
| JEV | 12/12 | 25.29<br>(0.31, 1.24) | 12/12 | 29.01<br>(0.17, 0.59) | 12/12 | 31.67<br>(0.16, 0.52) | 12/12 | 35.14<br>(0.41, 1.17) | 20/20 | 26.05<br>(0.83, 3.20) | 20/20 | 37.68<br>(0.63, 3.02) |

SD, standard deviation; %CV, percentage coefficient of variation; DENV, dengue virus; ZIKV, Zika virus; JEV, Japanese encephalitis virus; ND, not determined

Repeatability and reproducibility experiments evaluated the precision of the assay. The nucleic acids of the 6 arboviruses at concentrations of 1, 10, 100, and 1,000 copies/μL were used to assess the intra-assay repeatability in 12 replicates. The intra-assay repeatability results showed CVs below 2% for the detection of DENV1–4 and JEV at all dilutions. Similar results were seen for ZIKV at 100 and 1,000 copies/μL, but CVs were 4.67% for 1 copy/μL and 5.13% for 10 copies/μL. Inter-assay reproducibility was evaluated in 20 replicates at high (1,000 copies/μL) and low (100 copies/μL) concentrations across 5 days and between 2 operators. The assay also showed the CVs less than 5% in all viruses at both dilutions, ranging from 1.15%–3.20% for DENV4 and JEV detection at high concentration (1,000 copies/μL) **(Table 1)**

In addition, analytical specificity was evaluated in terms of cross reactivity using other pathogens, including Influenza A virus strain A/PR/8/34 (ATCC; VR-95), Influenza B virus strain B/Lee/40 (ATCC; VR-1535), RSV strain A-2 (ATCC; VR-1540), CHIKV strain ROSS, Enterovirus 71 (EV-71) strain H (ATCC; VR-1432), *Orientia tsutsugamushi* strain KARP (accession number = SAMN51290297), *Rickettsia typhi* strain SI-typh0423 (accession number = SAMN51290298). The assay showed no cross-reactivity with other pathogens **(Supplementary Table 6).**

The 150 archived clinical samples used to assess clinical performance for this assay, were comprised of 100 positive and 50 negative plasma samples for pan-flaviviruses collected during 2018-2024, were diagnosed at the time of collection by the commercial kits. The pathogen-specific real-time RT-qPCR–derived results from the designed primer-probe set and commercial kits were compared for individual samples **(Fig. 4C)**. The overall performance of the designed primer-probe was 97.33% accuracy (95% CI, 93.31–99.27%), 96% sensitivity (95% CI, 90.07% to 98.90%), and 100% specificity (95% CI, 92.89% to 100%) **(Table 2)**. For DENV1–4, our designed primer-probe showed 100% sensitivity (95% CI, 71.51 to 100%) and specificity (95% CI, 97.09 to 100%), while for ZIKV it showed 71.43% sensitivity (95% CI, 41.28 to 100%) and 100% specificity (95% CI, 97.32 to 100%) **(Table 2)**. Comparing the Ct value with the commercial kit, the overall Ct value from the designed primer-probe was significantly earlier to detect DENV1–4 (*p* = 0.0004, 0.0013, <0.0001, and <0.0001 for detection of the 4 DENV strains, respectively) but later to detect ZIKV (*p* = <0.0001) (**Fig. 4C)**. Bland-Altman analysis demonstrated strong concordance between 2 assays for DENV3 (mean ΔCt: -5.77, limit of agreement [LoA]: -8.12 to -3.42) **(Supplementary Fig. 3E)**, and ZIKV detection (mean ΔCt: 7.057, LoA: 3.10 to 11.01) **(Supplementary Fig. 3I)**. Most data points were distributed within the LoA, showing good consistency between the methods. However, only 4% (1 out of 25) of Ct values for DENV2, as well as 8% (2 out of 25) for DENV1 and DENV4, fell outside the 95% LoA (DENV1, -8.46 to 3.42 **[Supplementary Fig. 3A]**; DENV2, -7.39 to 3.39 **[Supplementary Fig. 3C]**; and DENV4, -5.99 to 0.396 **[Supplementary Fig. 3G]**). The correlation for pan-flaviviruses amplification obtained by 2 assays was moderate for DENV1 (r = 0.5486; *p* = 0.0045) **(Supplementary Fig. 3B)** and DENV2 (r = 0.6514; *p* = 0.0004) **(Supplementary Fig. 3D)**, while highly correlated for DENV3 (r = 0.9695; *p* = <0.0001) **(Supplementary Fig. 3F)**, DENV4 (r = 0.8114; *p* = <0.0001) **(Supplementary Fig. 3H)**, and ZIKV (r = 0.8081; *p* = 0.0047) **(Supplementary Fig. 3J)**.

**Table 2.** Clinical performance of the designed primers-probe.

| Pathogens | The number of samples |  |  | TP | FN | TN | FP | Sensitivity (%)<br>(95% CI) | Specificity (%)<br>(95% CI) | PPV (%)<br>(95% CI) | NPV (%)<br>(95% CI) | Accuracy (%)<br>(95% CI) |
| --- | --- | --- | --- | --- | --- | --- | --- | --- | --- | --- | --- | --- |
|  | Positive | Negative | Total |  |  |  |  |  |  |  |  |  |
| Pan-flavivirus | 100 | 50 | 150 | 96 | 4 | 50 | 0 | 96.00<br>(90.07-98.90) | 100<br>(92.89-100) | 100<br>(96.23-100) | 92.59<br>(82.71-97.03) | 97.33<br>(93.31-99.27) |
| - DENV1 | 25 | 125 | 150 | 25 | 0 | 125 | 0 | 100<br>(86.28-100) | 100<br>(97.09-100) | 100<br>(86.28-100) | 100<br>(97.09-100) | 100<br>(97.57-100) |
| - DENV2 | 25 | 125 | 150 | 25 | 0 | 125 | 0 | 100<br>(86.28-100) | 100<br>(97.09-100) | 100<br>(86.28-100) | 100<br>(97.09-100) | 100<br>(97.57-100) |
| - DENV3 | 11 | 139 | 150 | 11 | 0 | 139 | 0 | 100<br>(71.51-100) | 100<br>(97.38-100) | 100<br>(71.51-100) | 100<br>(97.38-100) | 100<br>(97.57-100) |
| - DENV4 | 25 | 125 | 150 | 25 | 0 | 125 | 0 | 100<br>(86.28-100) | 100<br>(97.09-100) | 100<br>(86.28-100) | 100<br>(97.09-100) | 100<br>(97.57-100) |
| - ZIKV | 14 | 136 | 150 | 10 | 4 | 136 | 0 | 71.43<br>(41.90-91.61) | 100<br>(97.32-100) | 100<br>(69.15-100) | 97.14<br>(93.69-98.73) | 97.33<br>(93.31-99.27) |

TP, true positive; FN, false negative; TN, true negative; FP, false positive; CI, confidence interval; PPV, positive predictive value; NPV, negative predictive value; DENV, dengue virus; ZIKV, Zika virus;

## Discussion

The global spread of *Orthoflavivirus* pathogens (such as DENV and ZIKV) presents a complex diagnostic challenge, as overlapping symptoms make clinical differentiation impossible. While RT-qPCR remains the standard for diagnosis, it is resource-intensive and struggles to detect rapidly mutating RNA viruses. The traditional approach focuses on designing primer-probe sets specific to individual species. This study proposes a systematic top-down AF computational framework that uses comparative genomic results using existing viral genomic data from public databases to produce a pan-viral genus detection assay. Our study adopted an AF k-mer strategy by decomposing genomes into fixed-length substrings (k-mers) and compared them with exact string statistics, unlike approximation results derived from biological sequence alignment. This approach treats the sequences as an unordered bag of features (k-mers), allowing for the rapid, unbiased processing of thousands of genomes to identify specific and conserved sequence “seeds” of the considered genomes for primer-probe design.

Analysis of 51 orthoflaviviral genomes revealed 3 regions of high conservation **(Fig. 2E),** including 5’ and 3’ untranslated regions (UTRs), which are replete with complex RNA secondary structures—stem-loops and dumbbells—that are vital for viral replication, cyclization, and host factor binding^41–43^. The 3’ UTRs are also the birthplace of subgenomic flaviviral RNA (sfRNA), a small, non-coding RNA produced when the host exoribonuclease (XRN1) degrades the viral genome in a 5’-to-3’ direction but stalls at the rigid pseudoknots of the 3’ UTR^44^. This results in the accumulation of 3’ UTR fragments at a level that can exceed that of genomic RNA by orders of magnitude^45^. While targeting the 3’ UTR can yield highly sensitive assays (due to the abundance of targets), it introduces biological noise. By choosing NS5 over the UTRs, our study opts for stoichiometric accuracy (measuring the actual virus genome) and broad inclusivity over the raw signal amplification offered by the sfRNA pool. This decision is central to understanding the sensitivity discrepancies observed in the clinical data, particularly for the Zika virus.

Our designed primer-probe set worked very well in the detection of 1–10 copies/µL of the standard virus references without cross-reactivity against non-target pathogens. The reproducibility is also high with %CV values less than 5.5% between assays, which is important when applying our assay in the routine lab settings. The assay in clinical samples achieved overall high accuracy and performance. Our designed primer-probe set performs better than the commercial kit for DENV serotypes, yielding significantly earlier Ct values. ZIKV detection by our assay showed less superiority to the commercial kit by later Ct values, which is a drawback of our assay. In addition, our assay missed detection of 4 archived ZIKV positive clinical samples, possibly due to a decay of viral material over more than 2 years of storage. However, using our primer-probe design set can substantially reduce assay costs for pan-viral species surveillance, through affordable custom synthesis.

In summary, we demonstrated the systematic top-down approach by an AF method for TaqMan RT-qPCR primer-probe design for pan-viral detection, enabling the ability to accurately detect multiple viruses in a considered *genus* rather than just the *species*. The workflow can be applied to other groups of viral pathogens of interest. In an interconnected world where the next pandemic threat is only a flight away, such broad-spectrum surveillance tools are the bedrock of global biosecurity.

## Author contribution (Contributor Role Taxonomy (CRediT))

Conceptualization: I.N., N.H. Data curation: K.S.

Formal analysis: K.S.

Funding acquisition: N.H., I.N.

Investigation: K.S., I.N., N.H., C.C., W.N., T.B, N.T.

Methodology: K.S., I.N., N.H., C.C

Project administration: N.T., N.H

Resources: N.H., N.T., C.C., W.N., T.B

Software: K.S., I.N., N.H

Supervision: I.N., N.H

Validation: C.C., N.H

Visualization: K.S., I.N.

Writing – original draft: K.S., N.H

Writing – review & editing: K.S., I.N., N.H.

## Competing interests

The authors declare no competing interests.

## Funding

This research was funded through a U.S. Centers for Disease Control and Prevention (CDC) cooperative agreement with the Thailand Ministry of Public Health entitled “THAI-GER: THAIland Genomic surveillance of Emerging infectious diseases facilitating Rapid response.” Cooperative agreement number 5U01GH002401-03. National Institutes of Health (P20GM125503) for partial support IN.

## Acknowledgements

We would like to extend our gratitude to Dr. R Suzanne Beard, and Dr. Pongpun Sawatwong from Division of Global Health Protection, CDC, Thailand for their contributions to project and validation design and for support through the cooperative agreement.

## References

1 Savić, V. et al. Zoonotic Orthoflaviviruses Related to Birds: A Literature Review. Microorganisms 13, 1590 (2025). 10.3390/microorganisms13071590

2 Haider, N. et al. Global dengue epidemic worsens with record 14 million cases and 9000 deaths reported in 2024. International Journal of Infectious Diseases 158, 107940 (2025). 10.1016/j.ijid.2025.107940

3 Zheng, J. et al. Global burden of dengue from 1990 to 2021: a systematic analysis from the Global Burden of Disease study 2021. Infectious Diseases of Poverty 14, 105 (2025). 10.1186/s40249-025-01365-x

4 Bhatt, S. et al. The global distribution and burden of dengue. Nature 496, 504–507 (2013). 10.1038/nature12060

5 Lim, A. et al. The overlapping global distribution of dengue, chikungunya, Zika and yellow fever. Nat Commun 16, 3418 (2025). 10.1038/s41467-025-58609-5

6 Quan, T. M., Thao, T. T. N., Duy, N. M., Nhat, T. M. & Clapham, H. Estimates of the global burden of Japanese encephalitis and the impact of vaccination from 2000-2015. Elife 9 (2020). 10.7554/eLife.51027

7 Anderson, J. F., Main, A. J., Cheng, G., Ferrandino, F. J. & Fikrig, E. Horizontal and vertical transmission of West Nile virus genotype NY99 by Culex salinarius and genotypes NY99 and WN02 by Culex tarsalis. Am J Trop Med Hyg 86, 134–139 (2012). 10.4269/ajtmh.2012.11-0473

8 WHO launches global strategic plan to fight rising dengue and other Aedes-borne arboviral diseases. Saudi Medical Journal 45, 1283–1284 (2024).

9 Chan, K. R. et al. Serological cross-reactivity among common flaviviruses. Frontiers in Cellular and Infection Microbiology 12, 975398 (2022). 10.3389/fcimb.2022.975398

10 Dias, B. D. P. et al. Challenges in Direct Detection of Flaviviruses: A Review. Pathogens 12, 643 (2023). 10.3390/pathogens12050643

11 Vina-Rodriguez, A. et al. A Novel Pan- *Flavivirus* Detection and Identification Assay Based on RT-qPCR and Microarray. BioMed Research International 2017, 1–12 (2017). 10.1155/2017/4248756

12 Madere, F. S. et al. Flavivirus infections and diagnostic challenges for dengue, West Nile and Zika Viruses. npj Viruses 3, 36 (2025). 10.1038/s44298-025-00114-z

13 Hill, V. et al. A new lineage nomenclature to aid genomic surveillance of dengue virus. PLoS Biol 22, e3002834 (2024). 10.1371/journal.pbio.3002834

14 Klontz, E. H. et al. Analysis of Powassan Virus Genome Sequences from Human Cases Reveals Substantial Genetic Diversity with Implications for Molecular Assay Development. Viruses 16 (2024). 10.3390/v16111653

15 Derelle, R. et al. Seamless, rapid, and accurate analyses of outbreak genomic data using split *k* - mer analysis. Genome Research 34, 1661–1673 (2024). 10.1101/gr.279449.124

16 Dong, Y., Chen, W.-H. & Zhao, X.-M. VirRep: a hybrid language representation learning framework for identifying viruses from human gut metagenomes. Genome Biology 25, 177 (2024). 10.1186/s13059-024-03320-9

17 Sankar, P., Sah, D., Kodati, D. & Dasari, C. M. Kmer-Based DNA Sequence Image Representation for Viral Disease, Translation and Mutated Pattern Prediction. BIO Web of Conferences 163, 01008 (2025). 10.1051/bioconf/202516301008

18 Brault, A. C., Fang, Y., Dannen, M., Anishchenko, M. & Reisen, W. K. A naturally occurring mutation within the probe-binding region compromises a molecular-based West Nile virus surveillance assay for mosquito pools (Diptera: Culicidae). J Med Entomol 49, 939–941 (2012). 10.1603/me11287

19 Alam, M. N. U. & Chowdhury, U. F. Short k-mer abundance profiles yield robust machine learning features and accurate classifiers for RNA viruses. PLOS ONE 15, e0239381 (2020). 10.1371/journal.pone.0239381

20 Ren, J. et al. Alignment-Free Sequence Analysis and Applications. Annual Review of Biomedical Data Science 1, 93–114 (2018). 10.1146/annurev-biodatasci-080917-013431

21 Ondov, B. D. et al. Mash: fast genome and metagenome distance estimation using MinHash. Genome Biology 17, 132 (2016). 10.1186/s13059-016-0997-x

22 Pierce, N. T., Irber, L., Reiter, T., Brooks, P. & Brown, C. T. Large-scale sequence comparisons with sourmash. F1000Res 8, 1006 (2019). 10.12688/f1000research.19675.1

23 Gardner, S. N., Slezak, T. & Hall, B. G. kSNP3.0: SNP detection and phylogenetic analysis of genomes without genome alignment or reference genome. Bioinformatics 31, 2877–2878 (2015). 10.1093/bioinformatics/btv271

24 Zielezinski, A., Vinga, S., Almeida, J. & Karlowski, W. M. Alignment-free sequence comparison: benefits, applications, and tools. Genome Biology 18, 186 (2017). 10.1186/s13059-017-1319-7

25 (NCBI), N. C. f. B. I. NCBI Virus, <https://www.ncbi.nlm.nih.gov/labs/virus/vssi/#/> (2004).

26 Shen, W. & Ren, H. TaxonKit: A practical and efficient NCBI taxonomy toolkit. Journal of Genetics and Genomics 48, 844–850 (2021). 10.1016/j.jgg.2021.03.006

27 Mihara, T. et al. Linking Virus Genomes with Host Taxonomy. Viruses 8, 66 (2016). 10.3390/v8030066

28 Pornputtapong, N. et al. KITSUNE: a tool for identifying empirically optimal k-mer length for alignment-free phylogenomic analysis. Frontiers in Bioengineering and Biotechnology Volume 8 - 2020 (2020). 10.3389/fbioe.2020.556413

29 Zhang, Q., Jun, S.-R., Leuze, M., Ussery, D. & Nookaew, I. Viral Phylogenomics Using an Alignment-Free Method: A Three-Step Approach to Determine Optimal Length of k-mer. Scientific Reports 7, 40712 (2017). 10.1038/srep40712

30 Marcais, G. & Kingsford, C. A fast, lock-free approach for efficient parallel counting of occurrences of k-mers. Bioinformatics 27, 764–770 (2011). 10.1093/bioinformatics/btr011

31 Do, D. T., Yang, M.-R., Vo, T. N. S., Le, N. Q. K. & Wu, Y.-W. Unitig-centered pan-genome machine learning approach for predicting antibiotic resistance and discovering novel resistance genes in bacterial strains. Computational and Structural Biotechnology Journal 23, 1864–1876 (2024). 10.1016/j.csbj.2024.04.035

32 Holley, G. & Melsted, P. Bifrost: highly parallel construction and indexing of colored and compacted de Bruijn graphs. Genome Biology 21, 249 (2020). 10.1186/s13059-020-02135-8

33 Langmead, B., Trapnell, C., Pop, M. & Salzberg, S. L. Ultrafast and memory-efficient alignment of short DNA sequences to the human genome. Genome Biology 10, R25 (2009). 10.1186/gb-2009-10-3-r25

34 Li, H. et al. The Sequence Alignment/Map format and SAMtools. Bioinformatics 25, 2078–2079 (2009). 10.1093/bioinformatics/btp352

35 Quinlan, A. R. & Hall, I. M. BEDTools: a flexible suite of utilities for comparing genomic features. Bioinformatics 26, 841–842 (2010). 10.1093/bioinformatics/btq033

36 Stadhouders, R. et al. The Effect of Primer-Template Mismatches on the Detection and Quantification of Nucleic Acids Using the 5′ Nuclease Assay. The Journal of Molecular Diagnostics 12, 109–117 (2010). 10.2353/jmoldx.2010.090035

37 You, Y. Design of LNA probes that improve mismatch discrimination. Nucleic Acids Research 34, e60–e60 (2006). 10.1093/nar/gkl175

38 Owczarzy, R. et al. IDT SciTools: a suite for analysis and design of nucleic acid oligomers. Nucleic Acids Research 36, W163–W169 (2008). 10.1093/nar/gkn198

39 Korkusol, A. et al. A novel flavivirus detected in two Aedes spp. collected near the demilitarized zone of the Republic of Korea. Journal of General Virology 98, 1122–1131 (2017). 10.1099/jgv.0.000746

40 De Oliveira Ribeiro, G., et al. Guapiaçu virus, a new insect-specific flavivirus isolated from two species of Aedes mosquitoes from Brazil. Scientific Reports 11, 4674 (2021). 10.1038/s41598-021-83879-6

41 de Borba, L. et al. RNA Structure Duplication in the Dengue Virus 3’ UTR: Redundancy or Host Specificity? mBio 10 (2019). 10.1128/mBio.02506-18

42 Liu, Y. et al. Structures and Functions of the 3’ Untranslated Regions of Positive-Sense Single-Stranded RNA Viruses Infecting Humans and Animals. Front Cell Infect Microbiol 10, 453 (2020). 10.3389/fcimb.2020.00453

43 Friebe, P. & Harris, E. Interplay of RNA elements in the dengue virus 5’ and 3’ ends required for viral RNA replication. J Virol 84, 6103–6118 (2010). 10.1128/JVI.02042-09

44 Besson, B., Overheul, G. J., Wolfinger, M. T., Junglen, S. & van Rij, R. P. Pan-flavivirus analysis reveals sfRNA-independent, 3’ UTR-biased siRNA production from an insect-specific flavivirus. J Virol 98, e0121524 (2024). 10.1128/jvi.01215-24

45 Pallares, H. M. et al. Zika Virus Subgenomic Flavivirus RNA Generation Requires Cooperativity between Duplicated RNA Structures That Are Essential for Productive Infection in Human Cells. J Virol 94 (2020). 10.1128/JVI.00343-20

